# Bacterial Community Shift in Marine Tar Balls following Low Salinity Exposure

**DOI:** 10.1101/241034

**Authors:** Mai Hoang Tran, Joong-Wook Park

## Abstract

The persistence and recurrence of tar balls causes detrimental effects on coastal wildlife, tourism, and fisheries. Previously, the human pathogen *Vibrio vulnificus* was detected in marine tar balls in the Gulf of Mexico. Marine tar balls also migrate to low-salt water bodies near the shore. The bacterial communities in tar balls from these environments, however, have not been fully characterized. Herein we describe our studies on the effect of reduced salinity on bacterial communities in marine tar balls. Tar balls collected from the Gulf of Mexico were incubated in deionized water for six months, and their microbial fingerprints were visualized using denaturing gradient gel electrophoresis. Our data show that the indigenous bacterial communities in marine tar balls shifted after being exposed to low-salinity condition. Dominant genera in the tar balls at low salinity were *Aquisalimonas* and *Parvibaculum*, neither of which includes known human pathogens.

## 1. Introduction

After being released into the ocean, petroleum hydrocarbons undergo multiple weathering processes, including evaporation, emulsification, dissolution, and oxidation [1]. Tar balls are a form of weathered oil occurring in discrete pieces of about 10 cm or less in diameter that can be washed onto beaches, float in water, or lie on the seafloor [1, 2]. Tar balls consist of a hard outer crust and a sticky semisolid core [3, 4]. The weathered oil in tar balls contains high concentrations of recalcitrant oil remnants such as long-chain alkanes (n>16), polycyclic aromatic hydrocarbons, and oxygenated hydrocarbons [5, 6]. In addition to the weathered tar balls, oil mats formed after oil spills under the ocean can be broken into small pieces by storms or hurricanes, producing less-weathered forms of tar balls [7]. Much effort has been made to clean the Gulf of Mexico after the Deepwater Horizon (DH) oil spill in 2010, but tar balls still reappeared after subsequent storms or hurricanes, causing recurrent problems along the Gulf coast.

The persistence of tar balls in ecosystems is a threat to coastal wildlife, tourism, and fisheries [8]. Recent research found that the DH-oil-spill-related tar balls contained stable free radicals and genotoxic metals such as chromium and nickel [9, 10], posing risks to human health. Additionally, a study by Tao *et al*. in 2011 showed that there was a 100-fold higher accumulation of *Vibrio vulnificus*, a human pathogen, in tar balls collected on the Gulf coast, compared to samples collected from sea water or sand [11]. This result suggests that the unique chemical and physical features of tar balls can create a niche for certain bacterial species, including human pathogens.

In October 2010, tar balls from the DH oil spill drifted to Lake Pontchartrain, an estuary on the Gulf coast [12, 13]. Changes in the ambient salinity as the tar balls traveled to the estuarine lake could affect the structures of their bacterial communities [14, 15]. Previous studies demonstrated that there were variations in microbial communities in water samples along a salinity gradient [16–18]. To the best of our knowledge, no research has been conducted on tar ball bacterial communities as they are exposed to low salinity environments. Given the fact that tar balls can nurture human pathogens [11], it is crucial to study the tar ball bacteria in estuarine environments. Here we investigated the bacterial community shift and dominant bacterial species in tar balls on exposure to low salinity condition.

## 2. Materials and Methods

### Sampling, microcosm preparation and DNA extraction

Tar ball samples were collected in September, 2012, immediately after Hurricane Isaac at Alabama beach (N30°14'30.23", W87°43'41.90") and stored at 4 °C for microcosm study and at -20 °C for initial bacterial community analysis. A microcosm was set up in October 2012 by placing tar balls with an equal volume of sterilized deionized water in a sterilized bottle. The tar balls were sampled after six months of incubation at 35 °C without shaking. The salinity of the microcosm was 495 ppm and the pH of the microcosm was 6.5. Total DNA was extracted from 0.5 g of tar ball samples using PowerSoil™ DNA Isolation Kit according to the manufacturer's instructions (MoBio Laboratories, Carlsbad, CA). Extracted DNA was kept at -20 °C prior to analysis.

### Polymerase chain reaction (PCR)

The primer set 341F with GC clamp and 534R was used to selectively amplify the V3 region of 16S rRNA gene from the total DNA [19]. One μL from each total DNA sample was mixed with 10 pmol of each primer, 0.25 mM of dNTP, 5 μL of 10x Green *Taq* PCR buffer, and 1 µL of Green *Taq* DNA polymerase to make a 50 µL PCR mixture (GenScript, Piscataway, NJ). PCR was set for initial denaturation at 95 °C for 5 minutes, followed by thirty five cycles, each comprising 20 seconds at 95 °C, 45 seconds at 55 °C, and 45 seconds at 72 °C. The reactions were finished with a final extension at 72 °C for 7 minutes. The amplification products were checked for integrity using 1.5% agarose gel electrophoresis.

### Denaturing gradient gel electrophoresis (DGGE)

Eight percent polyacrylamide gels were cast with a linear denaturing gradient of 40% on the top and 60% on the bottom. An equal volume of the PCR products from each sample was subjected to DGGE electrophoresis on a Bio-Rad DCode^TM^ system at 45 V for 16 hours in 1x Tris-acetate EDTA buffer (TAE, pH 8.0) (Bio-Rad Laboratories, Hercules, CA). Immediately after electrophoresis, the gels were stained with ethidium bromide and placed on a UV transilluminator for photographing (Fisher Scientific, Pittsburgh, PA).

### DGGE band excision and DNA sequencing

Selected DGGE bands were excised and submerged in 50 μL of deionized water for 24 hours. One μL of the supernatant containing the eluted DNA was used as template DNA for subsequent PCR-DGGE procedure (using the same experiment conditions as described above) to screen the purity of the excised DGGE band. When necessary, this process was repeated until one single DGGE band of interest was detected on the gels. The final PCR products were purified using MEGAquick-spin^TM^ Total Kit (iNtRON Biotechnology Inc., Korea) according to the manufacturer's instructions. The clean partial 16S rRNA gene amplicons were sequenced with forward and reverse primers separately by Genewiz Inc. (Genewiz Inc., South Plainfield, NJ). Each set of forward and reverse sequences was assembled into a consensus sequence using the DNA Subway program (Cold Spring Harbor Laboratory, Cold Spring Harbor, NY). The consensus sequences were aligned with 16S rRNA gene sequences in GenBank DNA libraries using Basic Local Alignment Search Tool (BLAST) [20].

### Gel Image Processing

Bionumerics version 7.6 platform was used to identify the bands and measure their intensity with respect to the background noise (Applied Maths, Austin, Texas).

## 3. Results and Discussion

Our data show that indigenous bacterial communities in marine tar balls shifted when exposed to a low salinity condition. Indigenous bacterial populations were decreased (bands A, B, and C), not changed (band D), or increased (band E) after 6 months of incubation in sterilized deionized (DI) water (Fig. 1). Since no microorganisms were introduced in the microcosm, all bacteria thriving in the tar ball samples were indigenous bacteria. Considering that the salinity of the Gulf of Mexico is 0 to 500 ppm in tidal fresh zones, 500 to 2,500 ppm in mixing zones, and above 2,500 ppm in seawater zones [21], our microcosms’ salinity of 495 ppm is categorized, for the purpose of this study, in the tidal fresh zone. A steep reduction in salinity from the seawater to the tidal fresh zone is considered to have triggered the bacterial community shift in the marine tar balls. Our observation was consistent with previous literature that showed different microbial diversity along salinity gradients in estuaries [17, 18] and a study reporting unique hydrocarbon-enriched bacterial communities established in response to changed salinity levels [16]. However, the studies mentioned above used water samples, while our research tested marine tar balls collected from the Gulf of Mexico.

**Figure 1.**
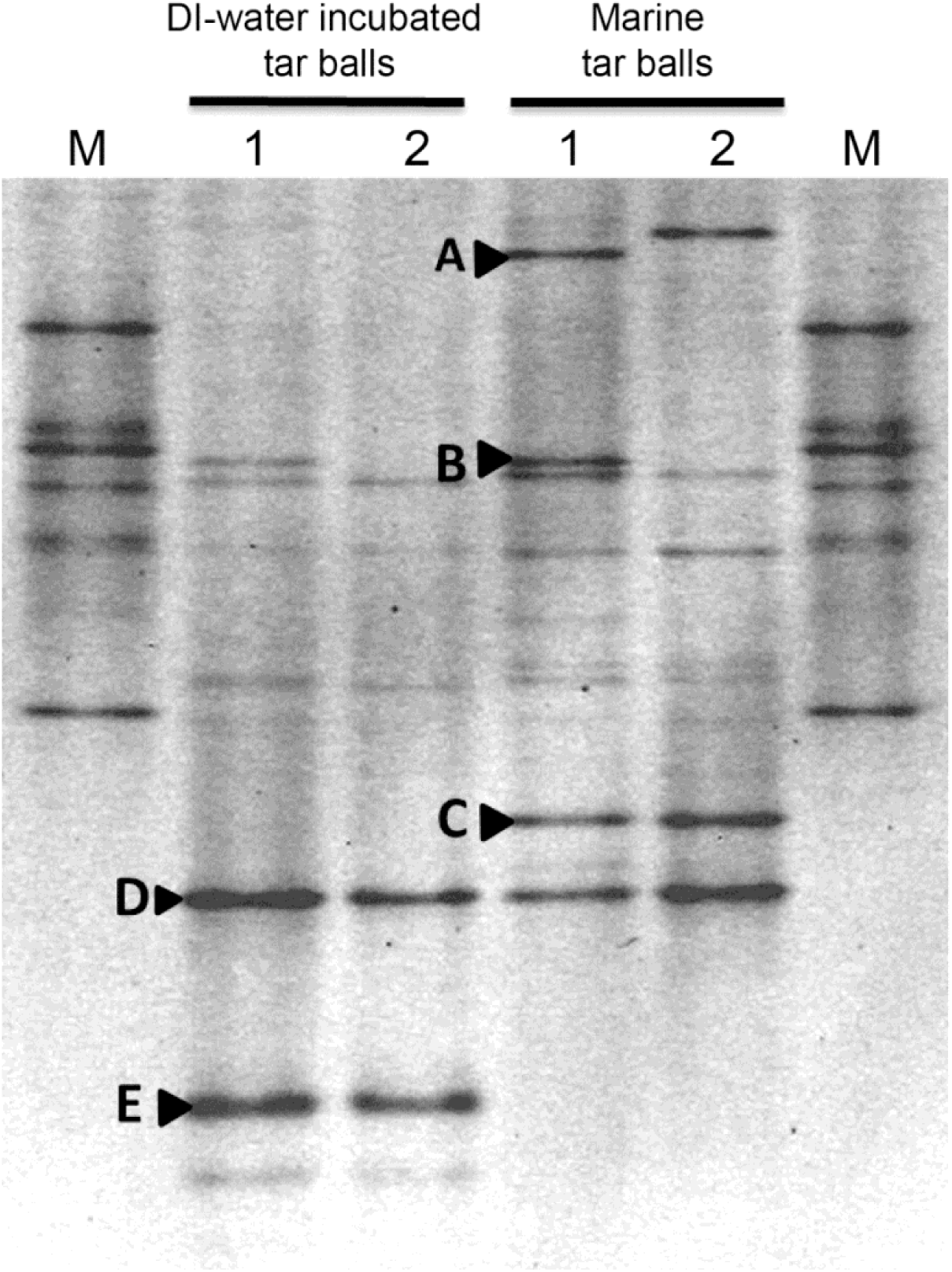
Bacterial community shift after exposure to low salinity conditions. Bacterial 16S rRNA gene profiles in untreated marine tar balls (Marine tar balls) and tar balls after six-month of incubation in deionized water (DI-water incubated tar balls) were analyzed by DGGE. Bands A to E were selected for sequencing. M: custom marker; numbers 1 and 2 are duplicates.

DGGE bands A, B, and C represent bacterial species dominant in marine tar balls, with their populations diminishing under low salinity conditions (Fig. 1). The DNA sequence of band A had an 85 *%* match in identity with the 16S rRNA gene of *Methylocystacceae bacterium* strain PKR-39 (Table 1), a type II methanotroph originally isolated from rhizosphere soil. Methanotrophic phylotypes were previously identified at the Gulf coast after the DH oil spill [22–24]. The presence of methanotrophs suggests hydrocarbons in the tar balls had been degraded into methane by methanogenic microorganisms [25, 26] when the tar balls were on the Gulf coast. DNA sequencing data of bands B and C showed that they belonged to members of the families *Sphingomonadaceae* (band B) and *Rhodobacteraceae* (band C), both of which are well-known marine taxa with an ability to degrade oil [27–29]. Multiple studies have found these taxa dominating in coastal water and sediments along the Gulf of Mexico after the DH oil spill [28–30, 32]. Moreover, *Rhodobacteracea* are known to be primary surface-colonizing bacteria in temperate coastal ecosystems, allowing other bacteria to colonize and grow on surfaces [33].

**Table 1.** Close relatives of partial 16S rRNA genes obtained from major DGGE gel bands

| Band | Close relative in GenBank databases | GenBank<br>Accession # | % Identity |
| --- | --- | --- | --- |
| A | *Methylocystaceae* bacterium PKR-39 | KJ000026 | 85 |
| B | *Novosphingobium panipatense*, isolate 1111TES1A3 | LN774374 | 90 |
| C | *Rhodobacterales* bacterium PSMR052 | KJ995712 | 96 |
| D | *Aquisalimonas halophila* YIM 95345 | KC577145 | 91 |
| E | *Parvibaculum* sp. MBNA2 | FN430653 | 97 |

Band D appeared in all tar ball samples with relatively consistent intensity (Fig. 1). The characterized genus closest to the 16S rRNA gene sequence in band D was *Aquisalimonas* (Table 1). Thus far, only three species of *Aquisalimonas* have been described - *A. asiatica, A. halophila*, and *A. lutea* - all of which are moderately halophilic and unable to survive in salinity concentrations lower than 1 *%* [34–36]. The dominance of band D under the low salinity condition suggests a novel *Aquisalimonas* species that thrive in salinity concentrations lower than 1 %, however, further study is necessary to confirm this observation. There has been no report on the pathogenicity of the three *Aquisalimonas* spp.

Band E was detected only in the tar ball samples incubated in sterilized DI-water for six months. As mentioned above, the bacterial species representing band E was indigenous to the marine tar balls, since no microorganisms were introduced in the microcosms. Therefore, it is likely that the abundance of species E in marine tar balls was below the detection limit of DGGE and increased under low salinity conditions [37]. DNA sequencing of this band showed that the species was closely related to *Parvibaculum* sp. MBNA2 (Table 1). This bacterial strain was previously isolated from freshwater sediments and a brackish water mud [38, 39]. Oiled wetland sediment and sea water samples collected in December 2010 along the Gulf of Mexico were reported to contain a large bacterial population affiliated with the genus *Parvibaculum*, [31]. Previous research has reported *Parvibaculum* spp. capable of degrading various hydrocarbon compounds, including PAH, alkanes, and surfactants [40–42]. No known human pathogens were reported belonging to the genus *Parvibaculum*.

In this research, we demonstrated that the indigenous bacterial community of marine tar balls was shifted when exposed to low salinity conditions and that dominant bacterial species in the tar balls at low salinity conditions were not phylogenetically close to potential human pathogens. The steep decline in salinity is most likely the major factor affecting the bacterial communities. However, this research did not consider other possible effectors during tar balls’ migration from the ocean to estuarine or inland water bodies, such as an introduction of exogenous bacteria and other environmental factors (e.g., temperature, pH, dissolved oxygen, and dissolved nutrients). We expect that our study provides preliminary data to support further research on pathogens in estuarine tar balls.

## Acknowledgements

We thank Dr. Ahjeong Son at Auburn University (currently at Ewha Womans University) for providing tar ball samples. This work was supported by the Chancellor's Fellowships and the Faculty Development Research grants, Troy University, Troy AL, and by the Beta Beta Beta Research Scholarship Foundation, Beta Beta Beta National Biological Honor Society.

